# Long-term Brillouin imaging of live cells with reduced absorption-mediated damage at 660nm wavelength

**DOI:** 10.1101/535419

**Authors:** Miloć Nikoliš, Giuliano Scarcelli

## Abstract

In Brillouin microscopy, absorption-induced photodamage of incident light is the primary limitation on signal-to-noise ratio in many practical scenarios. Here we show that 660 nm may represent an optimal wavelength for Brillouin microscopy as it offers minimal absorption-mediated photodamage at high Brillouin scattering efficiency and the possibility to use a pure and narrow laser line from solid-state lasing medium. We demonstrate that live cells are ~80 times less susceptible to the 660 nm incident light compared to 532 nm light, which overall allows Brillouin imaging of up to more than 30 times higher SNR. We show that this improvement enables Brillouin imaging of live biological samples with improved accuracy, higher speed and/or larger fields of views with denser sampling.

## 1. Introduction

Brillouin microscopy is an all-optical tool for measuring the mechanical properties of biological samples [1–3]. In recent years it has been applied in a number of studies of cells, tissues and biomaterials. Many comprehensive *in vivo* [4], and *ex vivo* [5] studies of the eye have been performed. Using Brillouin scattering, researchers have determined complete elastic tensor of a fibrous material [6]. The recent expansion of Brillouin microscopy into other biomedical fields has brought forward detailed studies of live non-labelled mammalian cells [7], plant cells and their environment [8], medically important tissue sections [9, 10], and protein concentrations in body fluids [11]. Most recently, Brillouin microscopy has been used to image whole mouse [12] and zebrafish embryos [13] as part of animal development studies.

Brillouin microscopy is based on confocally measuring the spontaneous scattering of light from thermal pressure/density waves inside a material [14–18]. Such spontaneous waves scatter the incident light and induce a frequency shift that is dependent on the local physical properties of the material. This provides an attractive solution to characterize material mechanical properties without contact and at high spatial resolution. However, the spontaneous Brillouin scattering signal is weak and the frequency shift associated with the scattering phenomenon is small — on the order of GHz [19]. This requires specialized spectrometers with high spectral dispersion and contrast; however these spectrometers have limited throughput [20]. As a result, high power of incident light or long exposure times must be used to obtain Brillouin spectra of high signal to noise ratio (SNR). In theory, this is not an issue, as the scattering process, unlike fluorescence, does not involve light absorption and thus it does not induce photodamage. However, within biological samples, in addition to being scattered, the incident light can also be absorbed by the endogenous chromophores of cells and tissues; and absorption can produce chemical changes or heat, and induce photodamage [21–24].

The dependence of absorption-induced damage on the wavelength and intensity of light is well characterized [22, 25]. Ideally, Brillouin scattering microscopy should be performed in the region of minimal absorption of biological cells and tissue, i.e. the so-called optical window in between the highly damaging UV and blue/green region where light can be absorbed by DNA, melanin, fat, bilirubin, or beta-carotene [24, 26], and the infrared region where water absorption becomes dominant. On the other hand, Brillouin scattering is a dipole-radiative process, thus the scattering efficiency is proportional to λ^−4^, so the signal intensity is significantly weaker at long wavelengths. An additional experimental consideration is that Brillouin microscopy requires stable (<GHz/hour), narrow (<MHz), and clean spectral lines (>80 dB). These specifications are met by gas lasers and solid-state lasers typically abundant in the blue/green region of the spectrum while semiconductor lasers, often used in the near-infrared region, usually present side modes and noise from amplified spontaneous emission. Indeed, Brillouin studies so far have used frequency doubled solid-state lasers, most using 532 nm wavelength of Nd:YAG laser, some using 561 nm [9, 27, 28] and one using 671 nm [29]. For *in vivo* studies, where photodamage is a strict concern, Brillouin studies have used near infrared wavelength (780 nm) employing semiconductor lasers and additional spectral purification elements [13, 30–32].

Here, we introduce Brillouin microscopy at 660 nm, which could represent an optimal compromise of fundamental and practical considerations: similar absorption profile as near-infrared 780 nm laser, but a significantly lower λ^−4^ penalty of the scattering cross section. In addition, the 660 nm line, like the widely adopted 532 nm line, can also be obtained from frequency doubling of Nd:YAG lasing transition, thus providing a clean and stable laser line for Brillouin measurements. Here, we characterize the Brillouin performance of this new wavelength on live cells in comparison to the 532 nm light. We demonstrate that the absorption-mediated damage to cells is dramatically reduced, thus increasing the range of power that can be delivered. We show that this significant improvement enables faster Brillouin measurements, higher precision spectral characterizations and/or long-term intracellular characterizations using Brillouin microscopy.

## 2. Methods

### 2.1 Cell Culture

Frozen NIH/3T3 (ATCC^®^ CRL1658™) cells were purchased from ATCC^®^ and cultured according to the supplier’s protocol. They were grown in T25 cell culture flasks in the standard 3T3 medium: Dulbecco’s modified Eagle medium (DMEM) supplemented with 4500 mg/L glucose, L-glutamine, and sodium bicarbonate, without sodium pyruvate (Sigma-Aldrich, catalog no. D5796). The DMEM formulation was supplemented with 10% (v/v) bovine calf serum (ATCC^®^ 30-2030™), and 50 U/mL penicillin and 50 μg/ml streptomycin (Thermo Fisher, catalog no. 15070063). The cells were grown at 37°C, in 5% CO_2_.

The cells were passaged according to the supplier protocol. First the cells were once washed with DPBS (Thermo Fisher). Next, 1 ml of 0.25% trypsin and 0.53mM EDTA solution (Sigma-Aldrich) was added to the cells and they were incubated at 37°C, in 5% CO_2_ for about 8 minutes until they were all detached from the T25 flask bottom. Next, they were covered with 9ml of the 3T3 medium and centrifuged at 150g for 5 minutes. The supernatant containing the trypsin-EDTA was removed, and the cells were resuspended in fresh 3T3 medium, and seeded into a new T25 flask. All the experiments were done on cells with passage number less than 10.

### 2.2 Preparation of cells for imaging

For Brillouin imaging and photodamage experiments we seeded the cells onto a glass bottom dish (Ibidi, μ-Dish 35 mm). Cells were seeded at low density, approximately 50,000 cells per 35mm plate. They were kept for at least 24 hours in the incubator to allow them to attach to the glass. The pH indicator commonly used in cell media is phenol red, which strongly absorbs visible light and can photosensitize cells [21]. Before the photodamage experiments, the cell medium was replaced with the cell medium without phenol red: 3T3 medium prepared as above, but with the phenol red free DMEM (ThermoFisher, catalog no. 21063029) instead of the regular DMEM. During the photodamage experiments cells were kept at 37°C using the incubation chamber for 35mm cell culture dishes (Warner Instruments). Brillouin images were acquired at room temperature. Prior to Brillouin imaging we allowed the glass bottom plate with cells on it to equilibrate to room temperature for 20 minutes.

To stain the cell nuclei, we used NucBlue™ Live ReadyProbes™ Reagent (Thermo Fisher, catalog no. R37605). NucBlue is a room-temperature stable, live cell stain formulation of cell permeable Hoechst 33342 nuclear dye that is excited at 360 nm and emits 460 nm fluorescence when bound to DNA. According to the protocol, we added 2 drops per ml of medium of NucBlue to cells plated on 35 mm glass bottom plate. The cells were incubated in the dark at room temperature for 15 min prior to imaging. Only cells that were used in Brillouin imaging experiments were stained.

### 2.3 Absorption-induced photodamage experiments

We coupled collimated laser light from a continuous-wave 532 nm laser (Torus, Laser Quantum, Inc.) directly into the Olympus 40x, 0.6 NA objective lens (Olympus, LUCPLFLN 40X) that was mounted on an inverted Olympus iX73 microscope. A variable neutral density (ND) filter (Thorlabs) was placed in the beam’s path before the objective to vary the intensity. Before placing the sample on the microscope stage, we recorded the power after the objective using the optical power meter (Thorlabs). Once the glass bottom plate was placed on the motorized microscope stage (Prior ProScan III), we moved the stage until the laser focal spot was inside an attached cell.

We imaged the laser-illuminated cell from the top side with a home-built Thorlabs cage system microscope: a 10x, 0.25 NA objective (Olympus) imaged onto the USB camera (Mightex Systems) with a spherical lens (Thorlabs). The sample was epi-illuminated with a red LED (Thorlabs, M660L4) using a polarizing beam splitter (Thorlabs) to separate the illumination and collection paths. We placed a 600 nm long pass filter (Thorlabs) after the 10x imaging objective to block the strong transmitted laser light that is focused in its imaging plane. Since the long pass filter makes the laser focal spot invisible in the camera image, and if it is not perfectly aligned it can translate the camera image, we first recorded the position of the laser focus in the image without the long pass filter. We found the position of the laser spot after the long pass filter insertion by simultaneously recording the position of a reference object at the top surface on the coverslip before and after the insertion of the long pass filter.

Photodamage experiments with 660 nm laser (Torus, Laser Quantum, Inc.) were performed on an inverted Olympus iX83 microscope with a motorized stage (Prior ProScan III). We illuminated the cells with a continuous wave collimated 660 nm laser light through the same objective (40x, 0.6NA) that we used in the 532 nm experiments. We used a notch filter that highly reflects (>98%) a narrow band of light at 660 nm (Chroma) to direct the laser light into the objective. This allowed us to image cell blebbing in transmission through the lower port of the microscope with a CCD camera (Mightex Systems). We put an additional short pass 600 nm filter (Thorlabs) before the camera to remove any laser light that would reach the camera due to the notch filter imperfection. We recorded the position of the laser focus in the image as described above.

To quantify photodamage, we used morphological changes in the cell, namely blebbing. Blebs form as the cell membrane integrity becomes compromised and the characteristic round shapes appear at the edge of the cell. Blebbing is a commonly used, reliable marker of photodamage [23, 25]. The time for a cell to develop blebs will be related to the total amount of energy absorbed by the cell during the illumination. Since we used focused light to irradiate cells, and all of the light was contained inside a cell, we related the total beam power (W) to the time until cell blebbed to get an estimate of the damage threshold of energy delivered for different wavelengths. Previous studies have reported light intensity (W/cm^2^) and energy density (J/cm^2^) in experiments where they used collimated light and illuminated a field of view much larger than a cell [22]. In our study, for the 532 nm photodamage experiments the range of total beam powers from was 2-50 mW used, and the corresponding peak intensity of light irradiated ranged from 1.78×10^5^ W/cm^2^ to 4.44×10^6^ W/cm^2^. The 1/e^2^ diameter of the Gaussian shaped laser beam entering the objective lens from the 532 nm laser was 1.8 mm. The focal spot size diameter was 1.69 μm. In our 660 nm photodamage experiments, the peak intensity of the center of the focal spot was 1.18×10^7^ W/cm^2^, corresponding to 40 mW total beam power after the objective. The beam size of the 660 nm laser entering the objective lens was 12 mm. The diameter of the focal spot was 0.93 μm with the Olympus 40x, 0.6 NA objective.

To acquire the time-lapse movies of cell reaction to light absorption, we used a home-built LabView program based on the acquisition software provided by the camera manufacturer. For all photodamage experiments, one brightfield frame was collected every 5 seconds until the cell showed visible blebs. The first frame was collected within 1 second of placing the cell in the focus of the laser. The frames were analyzed in ImageJ, and the time of the frame when the first bleb becomes visible was recorded.

### 2.4 Brillouin microscopy

To measure the Brillouin spectrum, we used an apodized two-stage VIPA spectrometer as described previously [7, 33]. The sample was illuminated with a linearly polarized 660 nm laser light through an overfilled 40x, 0.95 NA objective (Olympus, UPLSAPO40X2). Backscattered light was collected in the same objective and is coupled into a single mode optical fiber (Thorlabs, P1-630A-FC-2) with a fiber coupling lens (Olympus, 10x, 0.25 NA). Single mode optical fiber acted as a confocal pinhole and ensured that Brillouin microscope worked in a confocal configuration. The light from the fiber was coupled into the two-stage VIPA (15 GHz FSR, LightMachinery) spectrometer equipped with apodization [7], and coronagraphy [34] filters and imaged on the EMCCD camera (Andor iXon 897). We acquired the images by scanning the sample using the motorized stage (Prior ProScan III) controlled by the home-built NI LabView data acquisition program.

### 2.5 Brillouin spectrum analysis and image reconstruction

For each pixel of the image, we recorded one CCD camera frame. As described before [7], only the region of the camera with the anti-Stokes Brillouin peak and the Stokes Brillouin peak of the next diffraction order was recorded. We summed the pixel counts of 5 lines in the direction perpendicular to the spectral dispersion axis. We fitted each spectrum with Lorentzian peaks using least squares iterative method. The position of the peaks, their width and intensity were recorded, and a shift was calculated. The fitting and further analysis was done using MATLAB. We performed the calibration of pixel distances to frequency GHz by recording the Brillouin spectra of water and methanol, materials with known Brillouin shift [7].

To estimate the SNR and precision of a measurement we recorded 500 spectra of methanol. We calculated the signal by taking the average intensity of the maximum data point in the spectrum. The noise was calculated as the standard deviation of the same point. SNR is the ratio of the two. Precision was calculated after fitting and obtaining the shift value for each spectrum. Precision is the standard deviation of the 500 shifts that were measured. In Brillouin maps of the cells, we estimated the precision by calculating the standard deviation of a subset of pixels that measured the shift of the medium away from the cell.

### 2.6 Brightfield and confocal fluorescence imaging

We acquired brightfield images of the samples on the Olympus iX83 microscope in Köhler configuration. The cells were imaged in with the Olympus 40x, 0.95 NA objective (Olympus, UPLSAPO40X2), and with a CMOS camera (Andor Neo sCMOS). The fluorescence imaging was done with the same objective. We used Prior Scientific Lumen 200 light source to illuminate the sample in epi-fluorescence configuration. To image NucBlue stained nuclei of live cells, we used a standard DAPI fluorescence filter set for excitation at 350 nm and emission at 460 nm (Chroma). The emitted fluorescence was imaged with a spinning disk (Olympus IX2-DSU, spinning disk strip gap 18 μm) on the CMOS camera.

## 3. Results

### 3.1 Quantification of the absorption-induced cell damage

We quantified the photodamage of focused green (532nm) and red (660nm) light on the live fibroblast cells. We placed the NIH/3T3 cells (murine fibroblasts) in the focus of the laser light and recorded the time it takes for them to form blebs.

**Fig 1.**
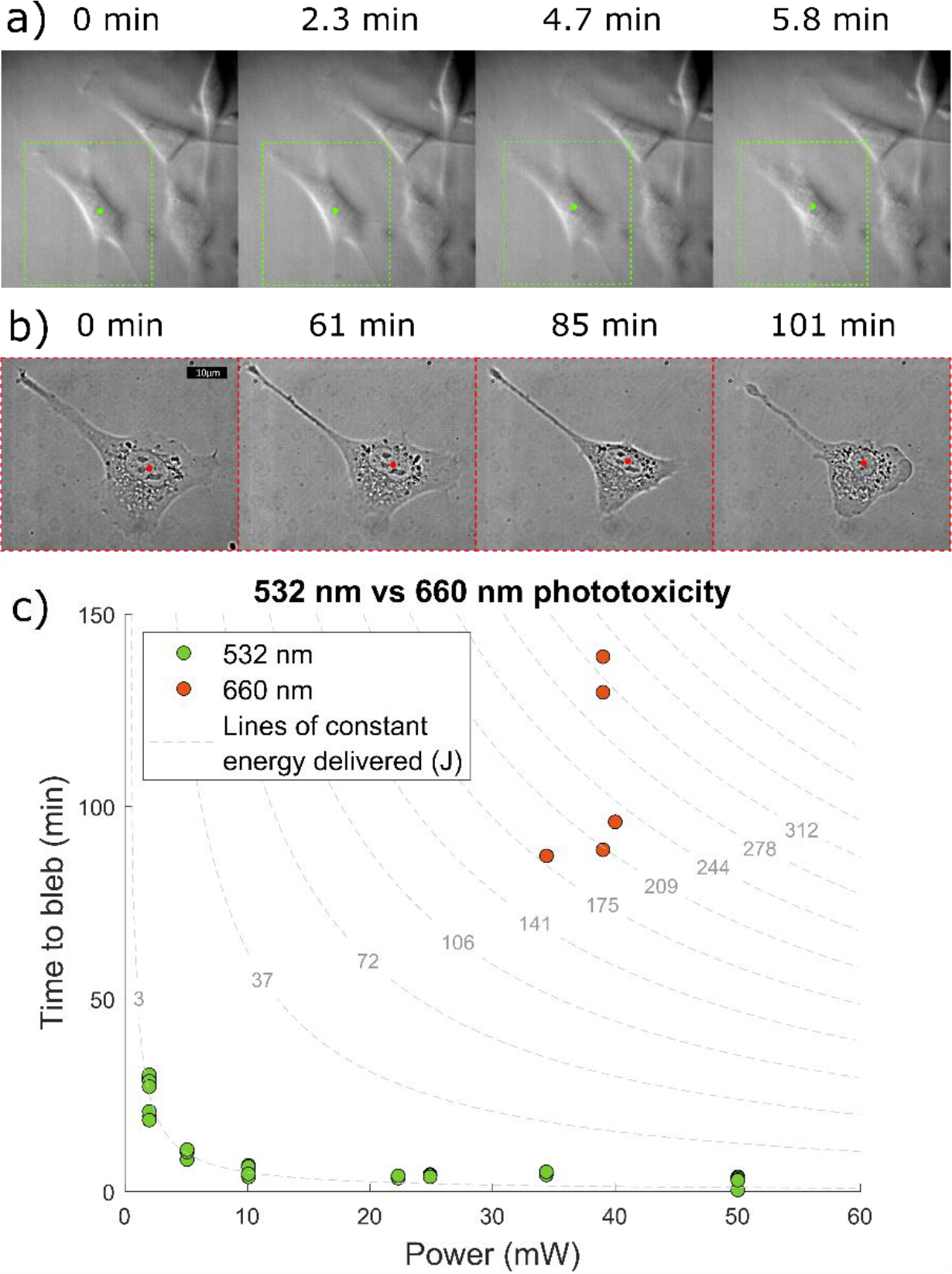
a-b) Representative frames of time-lapse movies that shows cell blebbing process when cells are exposed to the focus of a) 34 mW of 532 nm laser and b) 40 mW of 660 nm laser. The position of the laser focal spot is denoted by the dot. c) Measured time to bleb vs power delivered for green (532 nm) and red (660 nm) laser. Cell blebbing appears after a fixed amount of energy has been delivered to the cell in the 532 nm experiments. The points fall on the curve of 3.01 J delivered. The 660 nm experiments take a long time to show signs of damage. Because of cell variability and heat dissipation, blebbing times show a larger variation. Red cells need 2.49×10^2^ J on average to bleb.

As shown in Fig. 1, we observed that the cell appearance does not change significantly until a time when the blebbing process starts. At that time point, the cells contract their protrusions, and very soon the blebs start becoming visible. Only cells that are in the focus of the laser show this behavior.

We found that cells bleb within minutes of being exposed to the 532 nm light: ~5min for 34 mW. We varied the power delivered to the cells and found that, as expected, the time to bleb decreases proportionally with the increasing power, i.e. a constant amount energy needs to be delivered to the cells for the blebbing to occur. The cells take much longer time to form blebs when placed in the focus of the 660 nm light: >87 min for 40 mW. Due to the long experimental times, we only tested cells at the maximum power available in the setup with the 40x, 0.6NA objective. There is a larger spread of the blebbing times for red photodamage experiments, which could be attributed to the difficulty of maintaining consistent conditions over long experimental times (1-2 hours), i.e. there could be vertical drift of sample due to microscope instability, more heat dissipation within the cell, or variability between cells seeded on different plates.

On average, cells needed 249 J of irradiated energy at 660 nm to start forming blebs. The best fit constant energy curve to the data from the 532 nm experiment corresponds to the 3.01 J of energy delivered (R^2^=0.8918). Therefore, cells absorb the 660 nm light ~82 times less than the 532 nm. In other words, ~82 times more energy is needed to induce blebbing when cells are in the focus of the red laser.

### 3.2 Increased acquisition speed and spectrometer performance

To evaluate the performance of the 660nm-based spectrometer for Brillouin mapping of cells, we measured the signal to noise ratio (SNR) and the precision of the Brillouin frequency shift. We used the maximum power available in our setup: 61mW after the 40x, 0.95NA objective that is well suited for imaging single cells attached on glass in our setup [35]. 500 spectra of methanol were recorded at different exposure times. For each exposure time the SNR was quantified, as well as the precision of the estimated shift after fitting.

Fig. 2 shows that the SNR of the spectrometer is proportional to the square root of the illumination energy—higher exposure time at constant incident power. Remarkably, the red system provides shot noise limited spectra down to 0.3 ms of acquisition time. For shorter exposure times, the spectrum SNR becomes limited by the background camera noise.

**Fig 2.**
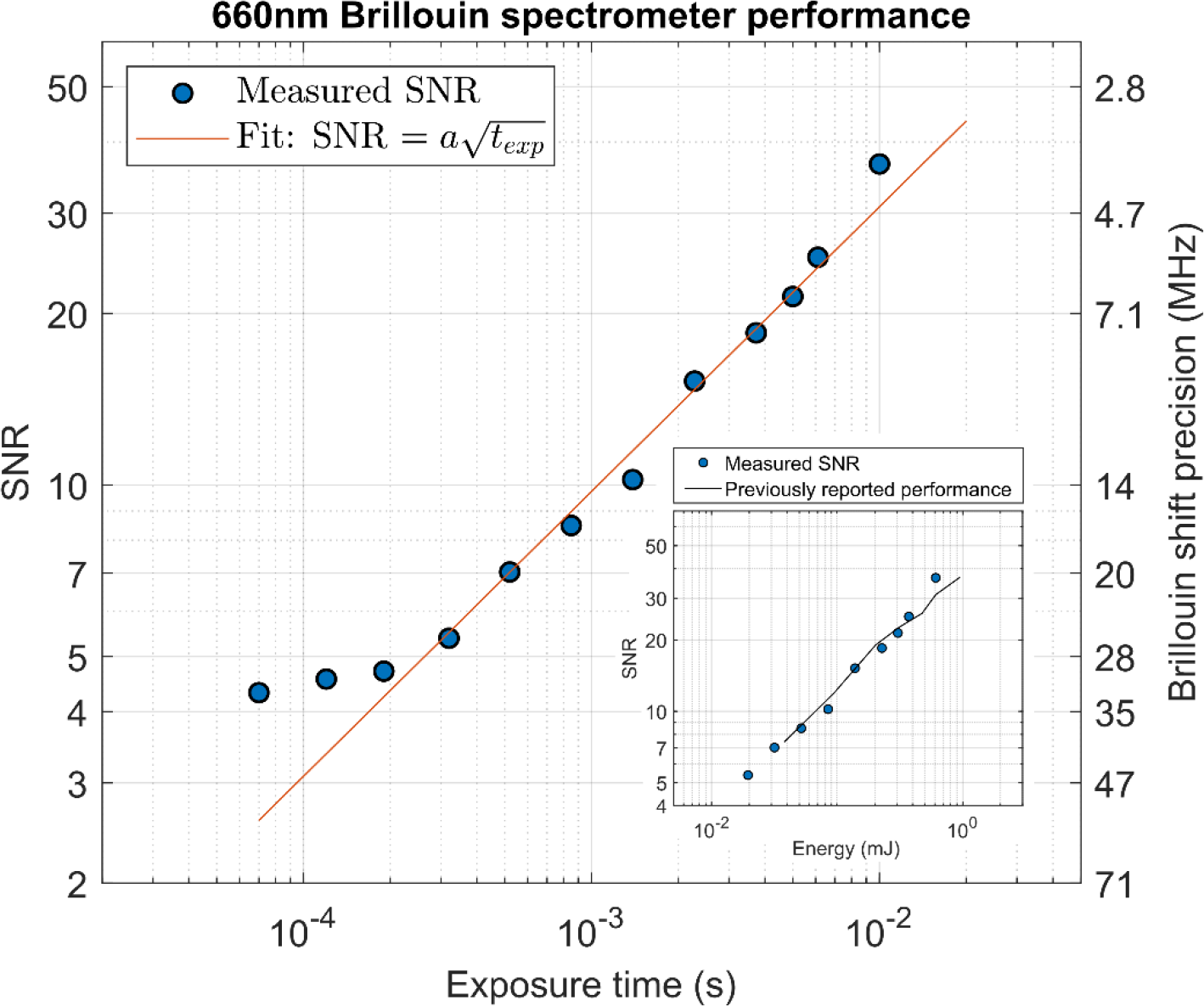
SNR and Brillouin shift precision vs the exposure time of methanol measured with 660 nm laser and 61 mW power. The setup is shot noise limited down to 0.3 ms, below which the camera noise becomes significant. The inset is SNR vs irradiated energy curve. Dark solid line is the previously published data from the 532 nm Brillouin spectrometer measuring methanol [7], rescaled by a factor of (532nm/660nm)^4^ to account for the lower scattering efficiency at 660 nm.

We also compared the performance of the spectrometer to the previously reported performance of equivalent 532nm-based spectrometers [7]. Since Brillouin scattering is proportional to the λ^−4^ we expect lower SNR for the same number of incident photons (or equivalently energy irradiated). In the shot noise limited region, we indeed see (532/660)^4^ = 42% of SNR when compared to the spectrometer performance at 532 nm. Therefore, our spectrometer performs equally to the previously reported values accounting for the lower scattering efficiency. Considering the fact that the red laser can deliver 82 times more energy to living cells without inducing damage, overall at 660 nm we can perform Brillouin imaging up to 34 times faster than before.

### 3.3 Improved Brillouin image quality due to improved precision in Brillouin shift

To illustrate the potential of improved precision, we imaged a 3T3 cell attached on a glass cover slip. We imaged the same part of the cell with two different integration times for each pixel imaged at 61 mW power. We estimate that the lateral resolution of the 40x, 0.95 NA at 660 nm is ~424 nm. The data in Fig. 3 was collected by imaging the part of the cell in steps of 83.3 nm with 20 ms exposure time. This was well below the optical resolution, which allowed us to average the Brillouin spectra from pixels in the local 3×3 neighborhood without introducing the blur artifact. Since we operated in the shot-noise limited regime the averaging of the signal from the 3×3 neighborhood is equivalent the signal that would have been obtained if we used 9 times longer exposure time and a pixel size of 250 nm. To obtain the image with pixel size of 250 nm and 20 ms exposure time, no averaging was performed and only the spectrum from the central pixel of the 3×3 neighborhood was used.

Fig. 3 shows the Brillouin image with 3.9 MHz precision, which corresponds to 0.064% relative error compared with an image of the same region with 9.2 MHz precision in Brillouin shift, which corresponds to 0.15% relative error, comparable to earlier reports [7]. Because of the power limitations of the setup, we achieved improved precision by increasing pixel exposure time. In the 660 nm experiment shown in the Fig. 3, the energy irradiated on the cell was ~5 times less than the energy needed to induce blebbing in the cells: (maximum 45 J delivered versus 250 J needed for blebbing).

The range of Brillouin shifts within a cell is on the order of 500 MHz. Nucleoli have only 50 MHz higher shift than their surroundings. To image them the precision should be at least one order of magnitude less than that difference. The improvement in precision is necessary to image tightly spaced intranuclear structure as shown in the Fig. 3. The 9.2 MHz error in the shift is high enough to affect the image quality and the ability to identify intracellular components. The ability to have more power, and image at comparable exposure times, allows us to clearly identify smaller and more densely spaced components inside the cell.

**Fig 3.**
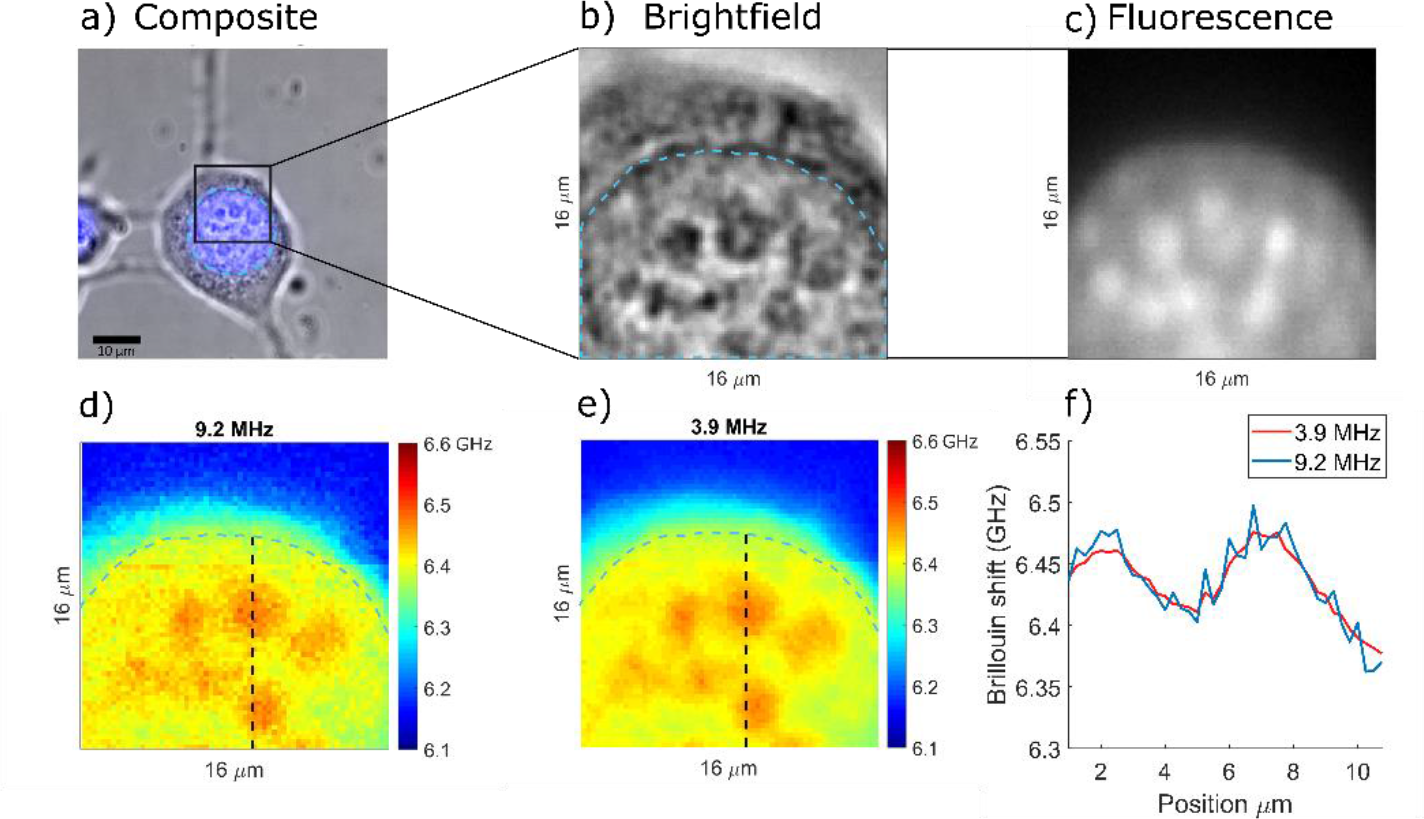
a) Brightfield and nuclear fluorescence composite image of a 3T3 cell attached on glass. b) Zoomed in region of the nucleus with clearly visible nucleoli. c) Spinning disk fluorescence image of the same region of the cell. d) Horizontal Brillouin shift confocal slice of a cell imaged with 9.2 MHz shift precision and 20 ms exposure time per pixel. d) Same horizontal Brillouin shift confocal slice imaged with 3.9MHz shift precision and 180 ms exposure time per pixel. Dashed blue line indicates the location of the nucleus edge. f) Line profile of the Brillouin images at two different exposure times. Location of the line profile is indicated by the black dashed line in the Brillouin maps in d) and e).

### 3.4 Long-term imaging, large field of view, and fast 3D mapping

With the higher laser power at 660nm, speed can be significantly improved without sacrificing the precision. This allows for much larger fields of view to be imaged. Often, the image acquisition time is the limiting factor in a Brillouin experiment. The cause for limited acquisition time could either be light induced damage, or a large number of experiments which limits the experimental time for measuring each cell or region of interest. If the total number of pixels that can be acquired is limited, experimenters either sacrifice the field of view, or spatial sampling (pixel size) regardless of the optical resolution provided by the objective used. In practice, a faster Brillouin spectrometer improves this tradeoff between the spatial sampling and the size of the field of view.

Here, we show a large field of view sampled at 500 nm step size. This pixel size is comparable to the optical resolution of the objective (424 nm). In the large field of view image, we are still able to clearly identify intranuclear structures. The improvement in speed allows for more room to acquire smaller pixels and avoid unnecessarily sacrificing the effective spatial resolution due to pixel size, even when imaging large fields of view.

**Fig 4.**
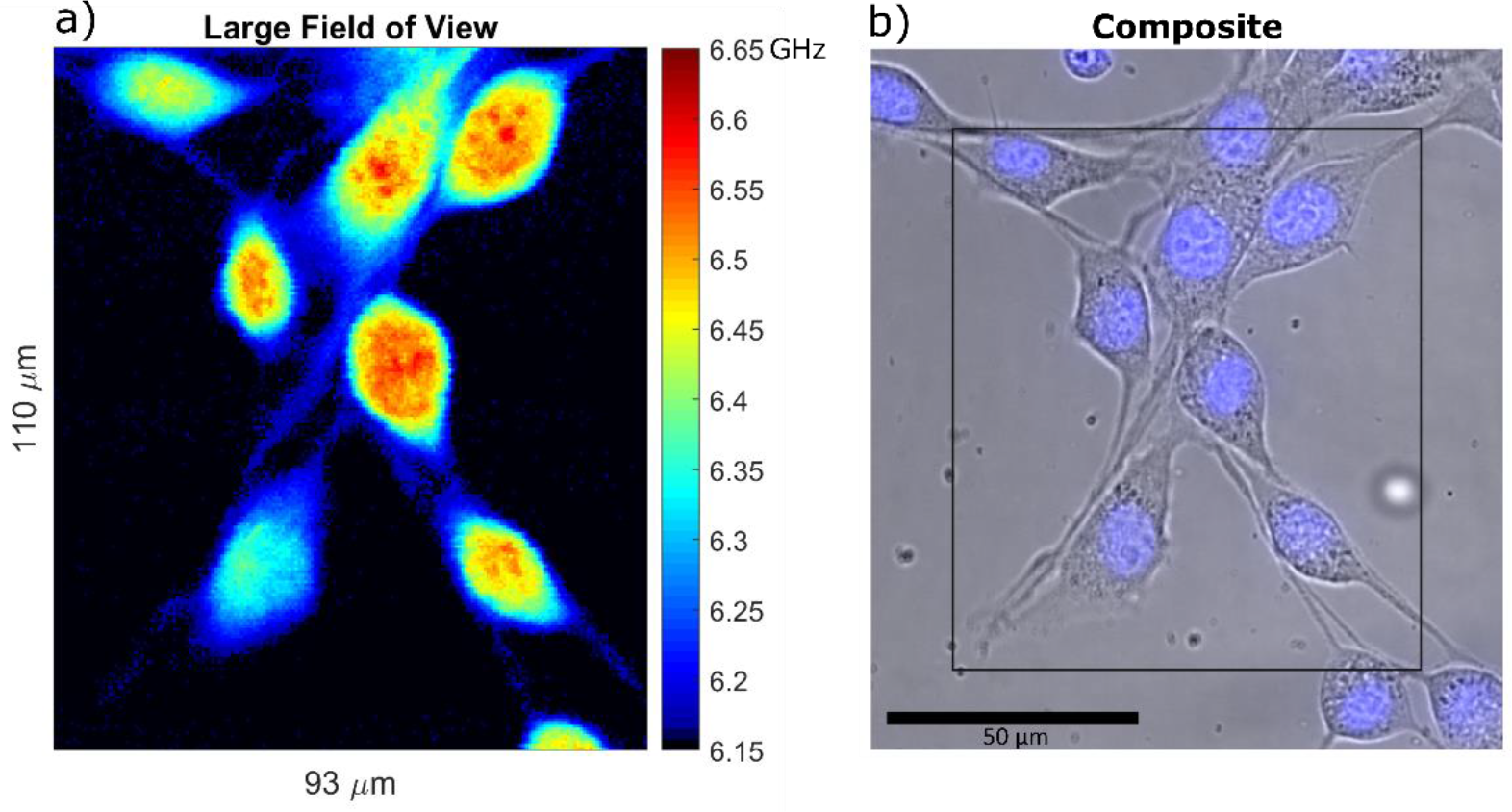
a) Example of a large field of view Brillouin shift image. The image is 110μm by 93μm, sampled at 0.5μm steps amounting to a total number of 40,920 pixels at 50ms exposure time, and total acquisition time of 34 min. Some of the cells in the image are only partly inside the horizontal confocal slice. Several nucleoli that fall within the confocal slice are visible. The lowest value of the colormap that corresponds to the measurement of cell medium is rendered black to better separate the cells from the background. b) Composite brightfield and nuclear fluorescence image of the scanned region.

To further illustrate the importance of higher limit on power allowed for Brillouin microscopy, we imaged a single attached cell in three dimensions with 1 μm 3D pixel size. Light induced damage is particularly limiting for tridimensional Brillouin imaging, since one cell or region of interest must be illuminated for a long time. In Fig. 5 we show that a complete Brillouin map of a NIH/3T3 cell on glass can be acquired within 7 min and 45 seconds. The cell was irradiated with at most 28.3 J of 660 nm light which is an order of magnitude less than the 250 J needed to induce blebbing.

**Fig 5.**
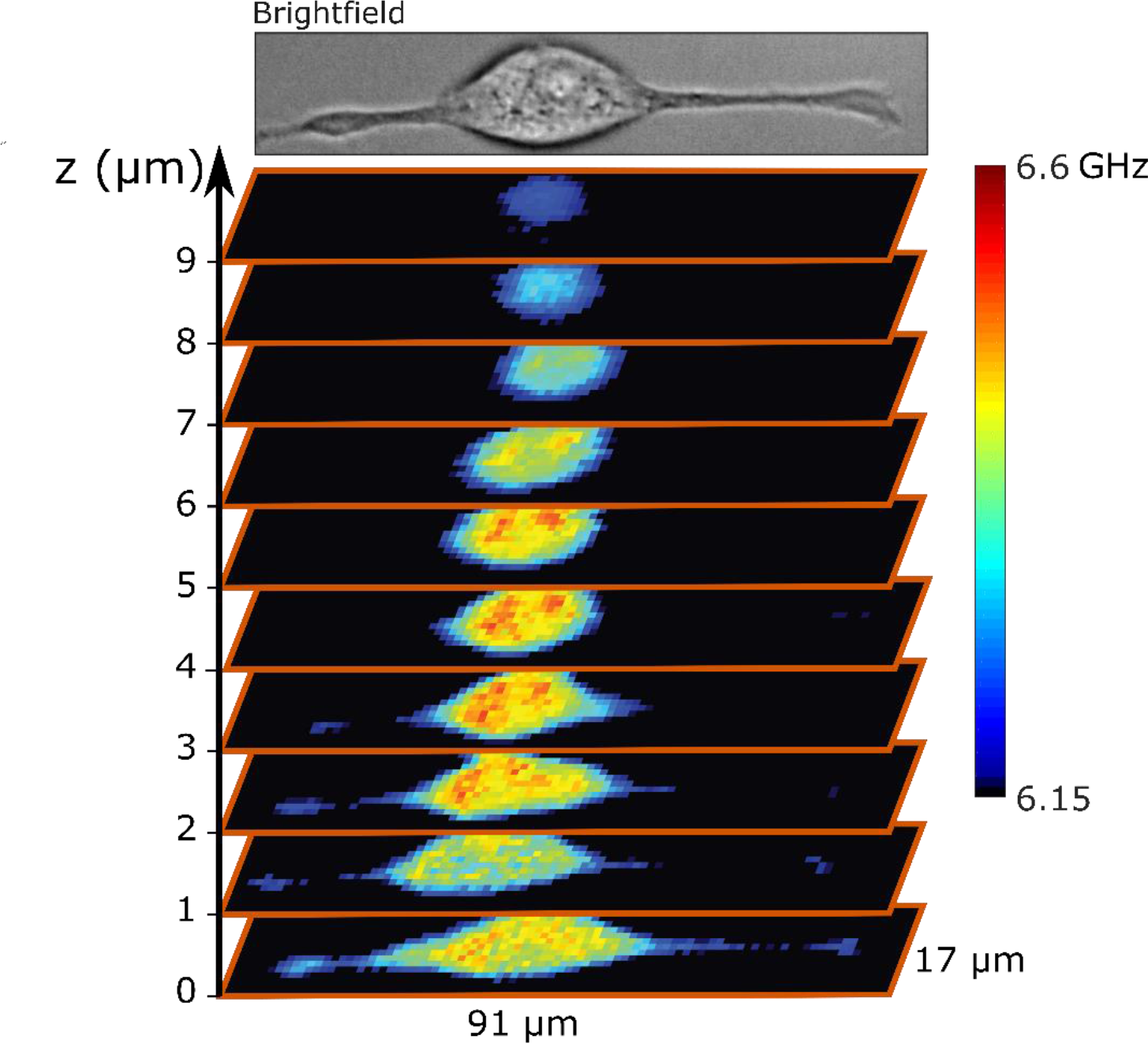
A 3D reconstruction of the Brillouin shift of a whole attached cell with the pixel size of 1μm×1μm×1μm. The whole acquisition took 7 min and 45 sec at 61 mW power. The lowest value of the colormap that corresponds to the measurement of cell medium is rendered black to better separate the cell from the background.

## 4. Discussion

Brillouin microscopy is based on spontaneous Brillouin light scattering, and due to weak scattering efficiency, it requires long integration times and high illumination powers. While scattering processes, like Brillouin scattering, do not induce thermal nor chemical changes in the sample, live cells and tissues can be damaged through the process of light absorption: either through the photochemical changes like creation of reactive oxygen species, or through heat dissipation produced by the absorption of the light [22, 23, 36].

The blue end of the visible light spectrum is very damaging to live biological samples. On the other hand, infrared light is strongly absorbed by water which could contribue to more intense light-induced heating. In choosing the optimal wavelength for Brillouin scattering, also the scattering cross section needs to be considered: due to the λ^−4^ dependence of the scattering efficiency, the Brillouin signal quickly decreases at longer wavelengths. Visible red light beyond 600 nm is minimally damaging to the cells, not significantly absorbed by water, and does not suffer from loss of scattering efficiency as much as previously used near-infrared wavelengths, thus it represents an optimal region of operation.

Here, we report the use of a clean and stable solid-state Nd:YAG laser at 660 nm that is in this optimal region of the electromagnetic spectrum for Brillouin imaging. Indeed, we found that one needs to irradiate 82 times more energy to induce the same amount of cell damage than with the previously used 532 nm. This is consistent with previous studies of light induced damage to live cells [22]. We show that after accounting for wavelength dependence of the scattering efficiency, the VIPA based spectrometer performs with the same quality as the previously reported 532 nm spectrometers. When lower scattering efficiency is considered together with the increased light induced damage limit on energy delivered, a VIPA based Brillouin spectrometer with a 660 nm laser can shorten the acquisition times ~34 times when compared to a 532 nm based one. We demonstrated that the reduced photodamage can be practically translated into long-term imaging, larger field of view, denser mapping, and better precision. The improvement in precision is particularly important for resolving those intracellular structures in living samples that are large enough to be imaged within diffraction limit and have been invisible to Brillouin microscopes so far due to the limited shift precision. An improvement in precision is also necessary for the ability to measure Brillouin linewidth, which is one order of magnitude smaller than the shift.

An additional advantage of the 660 nm illumination could materialize in experiments in which Brillouin maps are combined with fluorescence imaging [37, 38]. As fluorescent labels are a source of absorption and can significantly enhance the laser light damage to the live samples, care should be taken to avoid fluorescently labelling live samples with fluorescent dyes that have excitation near the Brillouin illumination wavelength. In this respect 660 nm is more suitable than shorter wavelengths because there is a wider variety of fluorophores that absorb in the blue/green range.

In this work, only the immediate and morphologically identifiable effects of light induced damage are addressed. This quantification of photodamage is appropriate for experimental scenarios when a single cell or tissue section is imaged at one time, and the measurements are not repeated on the same sample. This is relevant for experiments on individual live cells and organoids which can be seeded in large numbers, or for large tissue sections where each region is imaged only once, and many individual cells and regions can be imaged for each experimental condition. We did not observe any blebbing of neighboring non-illuminated cells in our samples. On the other hand, in experimental settings where long term effects of photodamage on live samples might be relevant, for example when individual cells are imaged at different time points, other methods for quantifying the damage might be more suitable [22, 23].

## Acknowledgments

The authors would like to thank Dr. Jitao Zhang and Antonio Fiore for useful discussions and a helping hand.

